# Surface versus volume synthesis governs growth-dependent efficacy of a *β*-lactam antibiotic

**DOI:** 10.1101/2024.01.31.578235

**Authors:** Rebecca Brouwers, Leonardo Mancini, Sharareh Tavaddod, Jacob Biboy, Marco Mauri, Elizabeth Tatham, Marie-Luise Enghardt, Ariane Zander, Pietro Cicuta, Waldemar Vollmer, Rosalind J. Allen

**Affiliations:** School of Physics and Astronomy, University of Edinburgh, James Clerk Maxwell Building, Peter Guthrie Tait Road, Edinburgh EH9 3FD, UK; Centre for Bacterial Cell Biology and Institute for Cell and Molecular Biosciences, Newcastle University, NE2 4AX Newcastle upon Tyne, UK; Biological and Soft Systems, Cavendish Laboratory, University of Cambridge, Cambridge, UK; Theoretical Microbial Ecology, Institute of Microbiology, Faculty of Biological Sciences, Friedrich Schiller University Jena, Jena, Germany; Cluster of Excellence Balance of the Microverse, Friedrich Schiller University Jena, Jena, Germany; Institute for Molecular Bioscience, The University of Queensland, Brisbane, Australia

## Abstract

The efficacy of *β*-lactam antibiotics depends strongly on bacterial growth rate. This can lead to poor correlation between in vivo action and in vitro assays, hindering effective prescribing – yet the mechanisms underlying growth-rate dependent *β*-lactam action remain unclear. Here, we investigate growth-rate dependent action of mecillinam, a *β*-lactam that targets the elongation-mediating PBP2 peptidoglycan transpeptidase enzyme, on *Escherichia coli* cells. We show that mecillinam alters the balance between the rates of cell surface area and volume synthesis in a growth-rate dependent manner. Under mecillinam treatment, cell volume increases exponentially at a rate fixed by the growth medium, but the cell’s ability to produce new surface area is compromised by the antibiotic. On rich medium, this imbalance leads to lysis, but on poor medium, slow-growing cells reach a new balance between surface area and volume synthesis, allowing sustained growth even at concentrations of mecillinam far above the EUCAST MIC value. A mathematical model based on surface area vs volume synthesis can quantitatively explain growth-medium dependent differences in mecillinam killing, as well as rescue from killing when cell morphology is perturbed in a microfluidic device. *β*-lactam antibiotic action is mechanistically complex, yet our work suggests that simple conceptual principles can help understand the interplay between molecular mechanism and cell physiology, potentially contributing to more effective use of these drugs.

## Introduction

The *β*-lactam group of antibiotics account for the majority of clinical antibiotic usage. It has long been known that these antibiotics are growth-rate dependent: slow-growing bacteria are less susceptible than fast-growing bacteria. Although this phenomenon was first reported for penicillin in the 1940’s and has been quantified in several studies (***Tuomanen et al., 1986; Cozens et al., 1986; Hadas et al., 1995; Lee et al., 2018)***, the underlying mechanisms remain poorly understood. Here, we provide a quantitative explanation for strongly contrasting efficacies of the *β*-lactam antibiotic mecillinam for fast and slow growing bacteria, based on the balance between surface and volume growth. Despite the molecular complexity of *β*-lactam antibiotic action, our work suggests that simple concepts may underly the growth-rate dependent efficacy of these antibiotics. *β*-lactam antibiotics interfere with synthesis of the peptidoglycan (PG) cell wall – a molecular mesh of glycan strands, connected by short peptides, that surrounds the cell membrane. Here we focus on the Gram negative bacterium *Escherichia coli*, which has a rod-shaped cell morphology and a thin (2-6 nm) PG layer, located in the periplasm. Since the PG mesh maintains both cell morphology and integrity in the face of turgor pressure, *β*-lactam exposure typically causes morphological changes and cell lysis (***Park and Strominger, 1957; Nonejuie et al., 2013; Kjeldsen et al., 2015; Yao et al., 2012)***. It is well established that the rate of lysis is proportional to growth rate pre-antibiotic exposure (***Tuomanen et al., 1986; Cozens et al., 1986; Hadas et al., 1995; Lee et al., 2018)***. However, the molecular mechanisms by which *β*-lactams cause lysis remain a topic of debate – changes in the balance between PG synthesis and hydrolysis (***Tomasz and Waks, 1975; Tomasz, 1979; Legaree et al., 2007)***, depletion of PG precursor molecules (***Banzhaf et al., 2012; Kohlrausch and Höltje, 1991)*** and futile synthesis-degradation cycling (***Cho et al., 2014)*** have been implicated.

The molecular targets of *β*-lactam antibiotics are the penicillin-binding proteins (PBPs), which incorporate nascent PG strands into the existing PG mesh (***Spratt, 1977)***. In *E. coli*, new PG incorporation is mediated by two dynamic complexes (***Typas et al., 2012)***. The elongasome mediates synthesis of the cylindrical cell wall during elongation; its key components are the TPase PBP2 and the GTase RodA associated with the actin-like cytoskeletal protein MreB. During cell division, PG synthesis for the new poles is carried out by a different complex, the divisome, containing the TPase PBP3, the GTase FtsW and the bifunctional TP-GTase PBP1b, ultimately controlled by the tubulin-like cytoskeletal protein FtsZ (***Egan and Vollmer, 2013)***. In addition to these complexes, *E. coli* also possesses other transpeptidase, transglycosylase and PG hydrolase enzymes involved with sacculus growth, maturation, remodelling and repair (***Egan et al., 2020; Morè et al., 2019; Peters et al., 2016)***.

Different *β*-lactam antibiotics target different TPase enzymes with varying degrees of specificity (***Spratt, 1977)***. Here, we focus on mecillinam, which binds specifically to PBP2, inhibiting cell elongation and causing cells to become round (***Lund and Tybring, 1972; Spratt, 1975; Jackson and Kropp, 1996)***. Introduced in 1972, mecillinam is effective for treatment of lower urinary tract infections (UTI).

PG metabolism is intimately linked with cell growth, since the PG layer needs to expand as the cell grows. Here, we connect *β*-lactam action with an emerging discussion about the balance between surface area and volume synthesis during cell growth. Recent studies have highlighted the close connection between cell shape and the relative rates of surface area and volume growth (***Harris and Theriot, 2016; Oldewurtel et al., 2021)*** – although the details of the mechanisms controlling surface area vs volume homeostasis are still emerging (***Shi et al., 2021; Oldewurtel et al., 2023)***. Other recent studies point to the combined roles of cell division control, surface area and volume synthesis in determining cell size and shape (***Basan et al., 2015; Lee et al., 2018; Ojkic et al., 2019; Serbanescu et al., 2020, 2021; Ojkic et al., 2022; Cylke et al., 2022; Howard et al., 2024)***. This discussion has implications for antibiotic action, since antibiotics may alter the balance between the rates of surface area and volume synthesis. Yet the nature of these implications remains somewhat unclear: some authors have suggested that changes in the surface area-volume synthesis balance could favour survival by reducing antibiotic influx (***Serbanescu et al., 2020; Ojkic et al., 2022; Cylke et al., 2022)***, but concomitant effects on nutrient intake (***Ojkic et al., 2022)*** and lysis rate (***Lee et al., 2018)*** may also be important.

In this work, we show that the balance between surface area and volume synthesis lies at the heart of the growth-rate dependence of mecillinam action. On rich media, fast-growing mecillinam-treated *E. coli* cells produce insufficient surface area relative to their rate of volume expansion, and they lyse. In contrast, for slow-growing cells on poor media, PBP2-independent divisome-mediated PG synthesis is sufficient to maintain surface area to volume balance, leading to sustained growth even at very high antibiotic concentration.

It is well known that antibiotic treatment strategies based on in vitro assays do not always correlate with in vivo therapeutic efficacy (***Cozens et al., 1986)*** – differences in microbial growth rate between the in vivo and in vitro environments are likely to be a contributing factor (***Zak and Sande, 1982; Cozens et al., 1986)***. Inaccurate prediction of in vivo efficacy leads to ineffective use of antibiotics, with associated risks of resistance emergence (***Albrich et al., 2004; Andersson and Hughes, 2014)***. This work contributes to our emerging understanding of how simple physiological principles can be used to explain complex, growth-dependent effects of antibiotics; these principles may ultimately help to achieve more effective and efficient antibiotic use.

## Results

### Mecillinam kills only fast-growing cells

To probe the interplay between bacterial growth rate and antibiotic action, we prepared cultures of *E. coli* in steady-state, exponential growth and exposed them to the *β*-lactam antibiotic mecillinam. Mecillinam binds to PBP2, inhibiting the attachment of the newly made PG chains to the sacculus during cell elongation (***Spratt, 1977; Cho et al., 2014)***. Comparing minimal media (MOPSgluMIN), media supplemented with amino acids (MOPSgluCAA) and media supplemented with amino acids and nucleotides (MOPSgluRDM) (see Methods) the antibiotic-free doubling time of our cultures varied almost by a factor of 2 (Fig. 1). To compare antibiotic inhibition dynamics on the three different media, we rescaled time by the antibiotic-free doubling time. In the absence of antibiotic, this caused the growth curves to collapse onto a single, media-independent, plot (Fig. 1a,b).

**Figure 1.**
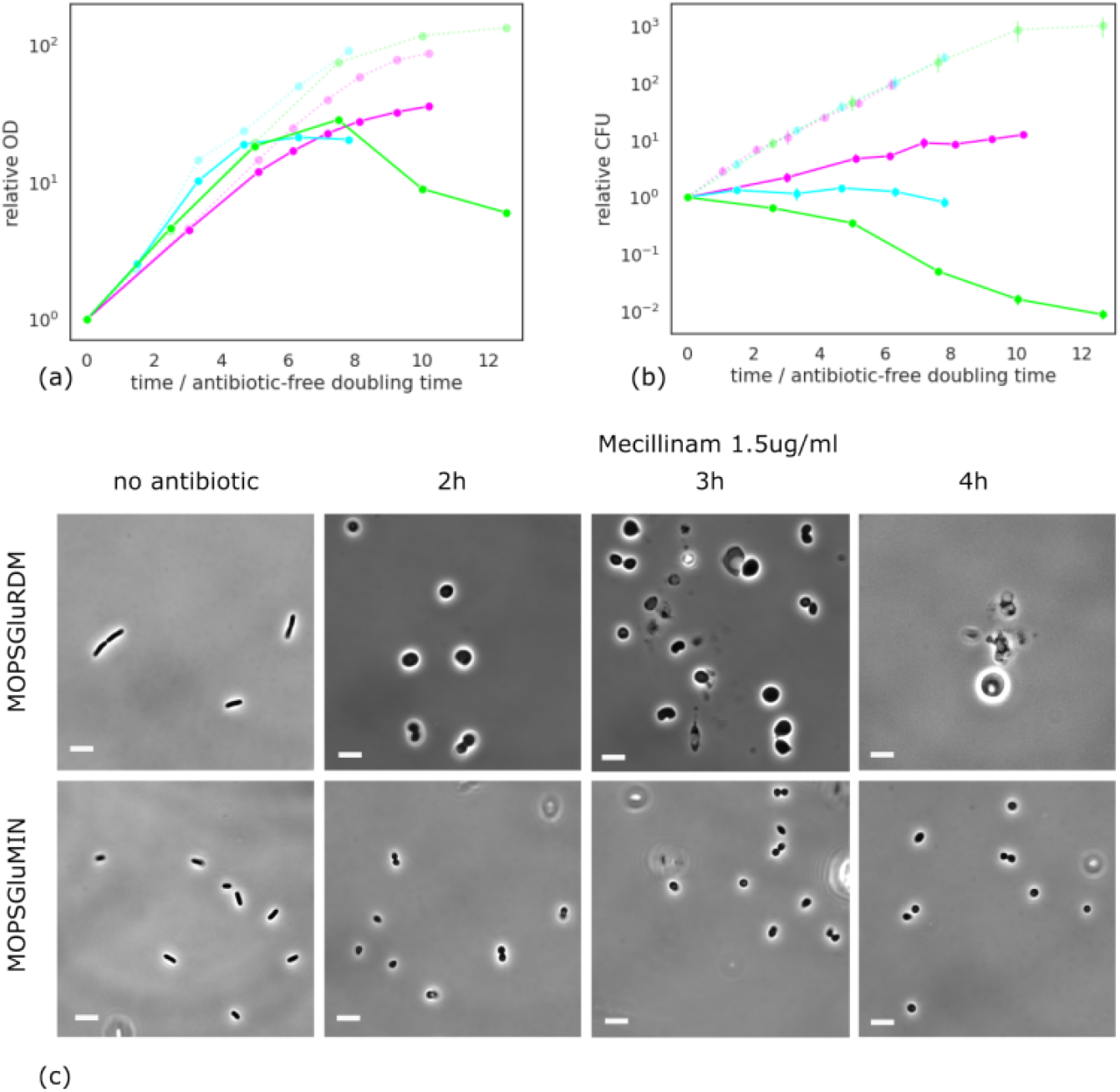
Contrasting efficacies of mecillinam for fast- and slow-growing bacteria. Steady-state exponential cultures of *E. coli* strain RJA002 were prepared on 3 media: MOPSGluRDM (data shown in lime-green), MOPSGluCAA (cyan) and MOPSGluMIN (magenta), at pH6.5, 34°C, and exposed to mecillinam at 1.5*μ*g/ml. (a): Optical density OD(600nm) of samples from the cultures, taken at different times after mecillinam addtion. OD is plotted relative to its value at time zero. Time is scaled by the doubling time of the no-antibiotic control cultures (29 ± 1min for MOPSGluRDM, 45 ± 4min for MOPSGluCAA, 55 ± 2min for MOPSGluMIN). The dashed lines show data for no-antibiotic controls; solid lines show mecillinam-treated cultures. (b): Live cell counts for samples from the same cultures, measured using colony counting (see Materials and Methods). (c): Phase contrast microscopy images of samples from the MOPSGluRDM and MOPSGluMIN cultures, taken at different times after mecillinam addition. On rich media (MOPSGluRDM) bacteria increase in size over a few cell cycles before lysing; on poor media bacteria proliferate as spheres in the presence of mecillinam. Scale bars are 5*μ*m in all images.

We exposed our cultures to mecillinam at a concentration of 1.5 *μ*g/ml. This corresponds to the MIC value of the parent strain MG1655 measured on LB media (Supplementary Methods), and is higher than the EUCAST MIC value of 1 *μ*g/ml (***EUCAST, 2020)***. After mecillinam was added, the optical density (OD) of our shake-flask cultures continued to increase at the pre-antibiotic rate for approximately 5 antibiotic-free doubling times (Fig. 1a). On rich media (MOPSgluRDM), the OD later decreased, indicating cell lysis, as previously reported (***Greenwood, 1976; Lee et al., 2018)***. For cells grown on minimal media (MOPSgluMIN) we observed no decrease in OD, suggesting that these cells did not lyse. Similar results were observed for higher and lower mecillinam concentrations.

Optical density measurements can be confounded by changes in cell size or shape (***Stevenson et al., 2016)***. Therefore, we also measured live cell numbers by colony counting (CFU; Fig 1b). Our CFU counts reveal that on rich media, cells start to die immediately after mecillinam is added even though the OD of the culture continues to increase (Fig 1a). In contrast, on poor media, CFU counts continue to increase exponentially after mecillinam addition, although at a decreased rate (Fig. 1b). CFU counts on the intermediate growth medium remained static after addition of mecillinam, suggesting a balance between killing and proliferation.

Phase-contrast microscopy of samples taken from our shake-flask cultures showed that mecillinam strongly affects cell morphology (Fig. 1c). *E. coli* cells become round upon exposure to mecillinam (***Lund and Tybring, 1972; Spratt, 1975; Nonejuie et al., 2013)***. Our data reveals that on rich media, the rounded bacteria increase dramatically in size, with some cell division events at early times, before lysing. On poor media, rounded bacteria remain small, and continue to divide (Fig. 1c). Our microscopy observations explain the apparent discrepancy between the dynamics of OD and CFU for cells exposed to mecillinam on rich media. At early times, cells become larger before lysing. Thus the total biomass (reflected in the OD (***Stevenson et al., 2016)***) increases in time, while the number of live cells decreases.

Taken together, our results point to strong growth-medium dependent effects of mecillinam on *E. coli*. On rich media, cells become round and divide a few times while increasing in size, and eventually lyse. In contrast, on poor media, cells become round but remain small and continue to successfully grow and divide, even at high concentrations of mecillinam.

### Changes in PG composition do not explain growth-medium dependence

Since mecillinam targets elongasome-mediated PG synthesis, we considered that the PG composition of cells might change under mecillinam treatment. To investigate if these changes could be growth-medium dependent, we performed HPLC analysis of muropeptides released from PG purified from cells that had been grown on rich media (MOPSgluRDM) vs poor media (MOPSgluMIN) and subjected to mecillinam treatment for 2 hours at 1.5 *μ*g/ml. As expected, our analysis revealed changes in PG composition for mecillinam-treated cells compared to the untreated control, in terms of glycan chain length, degree of cross-linking and tri- and tetra-peptide abundance. However, we did not observe substantial differences in PG composition (using the same metrics), for mecillinam-treated cells on rich vs poor media, apart from a longer glycan chain length present in cells grown in rich media (Table 1). This suggests that the medium-dependent response to mecillinam observed in our shake flas experiments (Fig. 1) was not caused by medium-dependent changes in PG composition.

**Table 1.** Analysis of PG composition. Muropeptide summary (%) or features for the PG isolated from RAJ002 grown in MOPSgluRDM and MOPSgluMIN media after exposure (or not) to 1.5*μ*g/ml mecillinam for 2h. Reported uncertainty refers to standard deviation of 2 replicate measurements.

|  | Feature of the PG from RJA002 cells grown in |  |  |  |
| --- | --- | --- | --- | --- |
|  | MOPSgluRDM | MOPSgluMIN | MOPSgluRDM | MOPSgluMIN |
|  | No antibiotic |  | Mecillinam |  |
| **Monomers (total)** | 58.0 ± 0.0 | 51.5 ± 1.2 | 49.2 ± 0.8 | 47.4 ± 1.6 |
| **Monomer di** | 2.0 ± 0.1 | 2.3 ± 0.1 | 3.2 ± 0.1 | 3.4 ± 0.0 |
| **Monomer tri** | 9.2 ± 0.4 | 9.1 ± 0.2 | 15.4 ± 0.7 | 13.5 ± 0.6 |
| **Monomer tetra** | 41.7 ± 1.4 | 36.5 ± 1.3 | 25.8 ± 0.6 | 26.4 ± 1.0 |
| **Monomer tetraGly4** | 1.1 ± 0.0 | 0.0 | 0.8 ± 0.0 | 0.0 |
| **Monomer penta** | 0.0 | 0.0 | 0.0 | 0.0 |
| **Monomer anhydro** | 0.0 | 0.3 ± 0.1 | 0.2 ± 0.3 | 0.5 ± 0.1 |
| **Dimers (total)** | 37.4 ± 1.7 | 39.5 ± 0.3 | 39.3 ± 0.7 | 37.6 ± 0.3 |
| **Dimers(DD)** | 35.9 ± 2.2 | 37.5 ± 0.3 | 36.2 ± 0.4 | 34.5 ± 0.4 |
| **Dimers(LD)** | 1.4 ± 0.3 | 1.8 ± 0.2 | 2.5 ± 0.3 | 2.7 ± 0.0 |
| **Dimers anhydro** | 1.3 ± 0.2 | 2.9 ± 0.5 | 3.2 ± 0.5 | 3.5 ± 0.1 |
| **Trimers (Total)** | 2.7 ± 0.5 | 6.0 ± 0.2 | 6.8 ± 0.7 | 8.6 ± 0.7 |
| **Trimer (anhydro)** | 0.2 ± 0.3 | 2.0 ± 0.1 | 2.2 ± 0.5 | 3.4 ± 0.7 |
| **Tetramers(Total)** | 0.0 | 1.1 ± 0.7 | 0.8 ± 1.2 | 2.1 ± 1.1 |
| **Tetramer (anhydro)** | 0.0 | 1.1 ± 0.7 | 0.8 ± 1.2 | 2.1 ± 1.1 |
| **Dipeptides (total)** | 2.0 ± 0.1 | 2.3 ± 0.1 | 3.2 ± 0.1 | 3.4 ± 0.0 |
| **Tripeptides (total)** | 12.7 ± 0.2 | 14.7 ± 0.3 | 25.5 ± 0.3 | 22.4 ± 0.3 |
| **Tetrapeptides (total)** | 80.3 ± 2.6 | 77.6 ± 1.1 | 63.6 ± 2.1 | 65.8 ± 0.2 |
| **Pentapeptides (total)** | 0.3 ± 0.0 | 0.4 ± 0.0 | 0.3 ± 0.0 | 0.4 ± 0.0 |
| **Lpp Attachment (%)** | 4.8 ± 2.5 | 4.6 ± 0.4 | 5.9 ± 1.9 | 6.3 ± 0.0 |
| **LD-cross-links (%)** | 0.7 ± 0.2 | 0.9 ± 0.1 | 1.3 ± 0.1 | 1.3 ± 0.0 |
| **DD-cross-links (%)** | 19.8 ± 0.8 | 23.6 ± 0.5 | 23.2 ± 1.6 | 24.6 ± 1.1 |
| **Chain ends (%)** | 0.7 ± 0.0 | 2.4 ± 0.4 | 2.5 ± 0.7 | 3.4 ± 0.5 |
| **Average glycan chain length (DS units)** | 138.9 ± 1.4 | 42.4 ± 7.7 | 41.3 ± 11.6 | 29.5 ± 4.6 |
| **Degree of cross-linkage** | 20.5 ± 0.5 | 24.6 ± 0.5 | 24.8 ± 1.7 | 26.2 ± 1.1 |
| **% peptides in cross-links** | 42.0 ± 0.0 | 48.5 ± 1.2 | 50.8 ± 0.8 | 52.6 ± 1.6 |

### Mecillinam changes the balance between area and volume synthesis

Since mecillinam inhibits the incorporation of new PG in the side wall of the rod-shaped cells, we hypothesized that the balance between volume and surface area synthesis might be changed upon exposure to mecillinam. To investigate this, we used a microfluidic mother machine device (***Panlilio et al., 2021; Wang et al., 2010; Long et al., 2013)*** to measure rates of volume and surface area growth for individual cells before and after exposure to mecillinam. Bacteria were grown in microfluidic channels for 10 h in either rich or poor media (MOPSgluRDM or MOPSgluMIN) before mecillinam was added at 1.5 *μ*g/ml (Materials and Methods). In this device, mecillinam exposure causes cells to swell laterally but they remain confined within the channel, so that their length and width can be tracked. Inferring from this data the rates of surface area and volume growth (Materials and Methods), we observed that the rate of volume growth did not change substantially upon addition of mecillinam, on either rich or poor media. However, the rate of surface area growth decreased after the antibiotic was added (Fig. 2). Therefore the balance between the rates of volume and area synthesis is changed in the presence of mecillinam.

**Figure 2.**
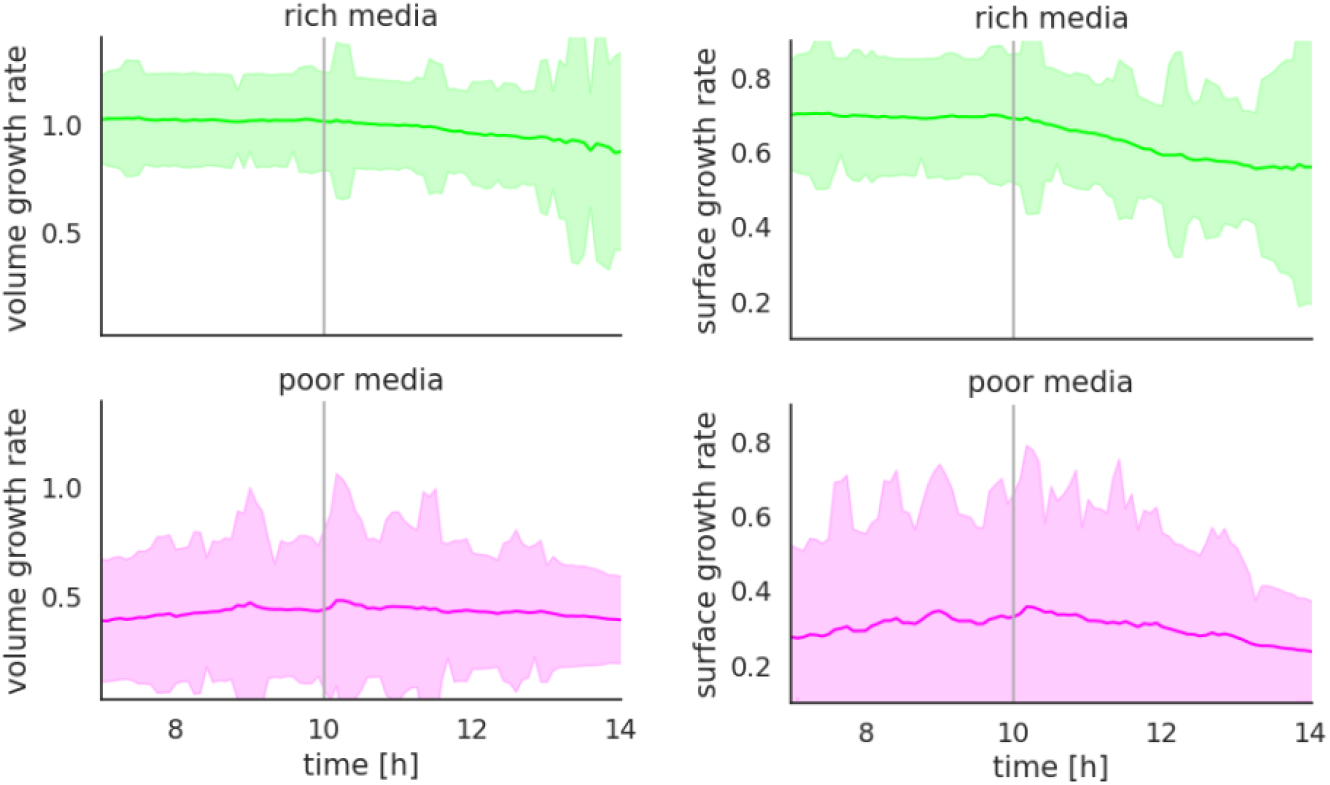
Microfluidic measurement of volume and surface area growth rates. The average volume and surface area growth rates for bacterial cells confined in channels in a microfluidic mother machine are shown as a function of time before and after addition of mecillinam at 1.5*μ*g/ml. Volume and surface area growth rates are inferred from analysis of cell length and width dynamics (see Material and Methods). The addition of mecillinam at 10h is indicated by the vertical line. Data is shown for rich media (MOPSGluRDM, pH6.5, 24°C) and poor media (MOPSGluMIN, pH6.5, 34°C).

### Area-volume model predicts growth-rate threshold for efficacy of mecillinam

Our data show that *E. coli* cells on rich media become round and increase in size over several cell cycles, before lysing, while cells on poor media also become round, but continue to grow and divide, maintaining a stable, smaller, cell size (Fig. 1). To rationalize these observations, we propose a model in which mecillinam (i) causes cells to become spherical (consistent with Fig. 1), and (ii) decreases the rate of surface area synthesis, but not the rate of volume synthesis (consistent with Fig. 2).

In this model, since the rate of volume synthesis is unperturbed by antibiotic, the cell doubles its volume in time *T*_vol_ = *T*_0_ (the cell cycle time in the absence of antibiotic, which is set by the growth medium; *T*_0_ = 2*π*/*λ*_0_ where *λ*_0_ is the antibiotic-free growth rate). To achieve a stable cell cycle, the cell needs to match its rates of surface area and volume synthesis; in other words, the surface area must double in time *T*_0_. For a sphere of radius *R*, the surface area is 4*πR*^2^. This implies that, to achieve a stable cell cycle with a birth radius *R*, the cell has to maintain a surface area synthesis rate (averaged over the cell cycle) of *Y*_t_ = 4*πR*^2^/*T*_0_. If we assume that a cell immediately rounds up on exposure to mecillinam, its initial radius *R* can be related to the antibiotic-free volume at birth *V*_0_by *R* = 3*V* ^1/3^/(4*π*). The threshold area synthesis rate for stable growth is then *Y*_t_ = 9*V* ^2/3^/(4*πT*_0_).

If the area synthesis rate falls below the threshold *Y*_t_, the surface area to volume ratio must decrease, which implies that the cell must become larger (since for a sphere, surface area/volume scales inversely with *R*). Our model therefore predicts that an insufficient rate of surface area synthesis relative to volume synthesis will cause the cell to increase in size over successive cell cycles (by taking a longer time to complete division), until, we suppose, it becomes too large to be able to divide, and lyses.

This prediction is consistent with the growth-rate dependent response to mecillinam observed in our experiments. Using data from ***Taheri-Araghi et al.(2015)*** for the birth volume of *E. coli* cells during exponential growth for a range of exponential growth rates, we plotted the model prediction for the threshold surface area synthesis rate *Y*_*t*_ that is required for continued proliferation in the presence of mecillinam (Fig. 3). The value of *Y*_*t*_ decreases steeply as the doubling time increases. Superimposing our mother machine data for the surface area synthesis rate for cells exposed to mecillinam in narrow channels (Fig. 2), averaged over a 1h timeframe after addition of mecillinam, we find that on the rich media (MOPSgluRDM) the rate of surface area synthesis is below the predicted threshold, while on the poor media (MOPSgluMIN) it is above the threshold. Therefore our model is quantitatively consistent with our experimental observations.

**Figure 3.**
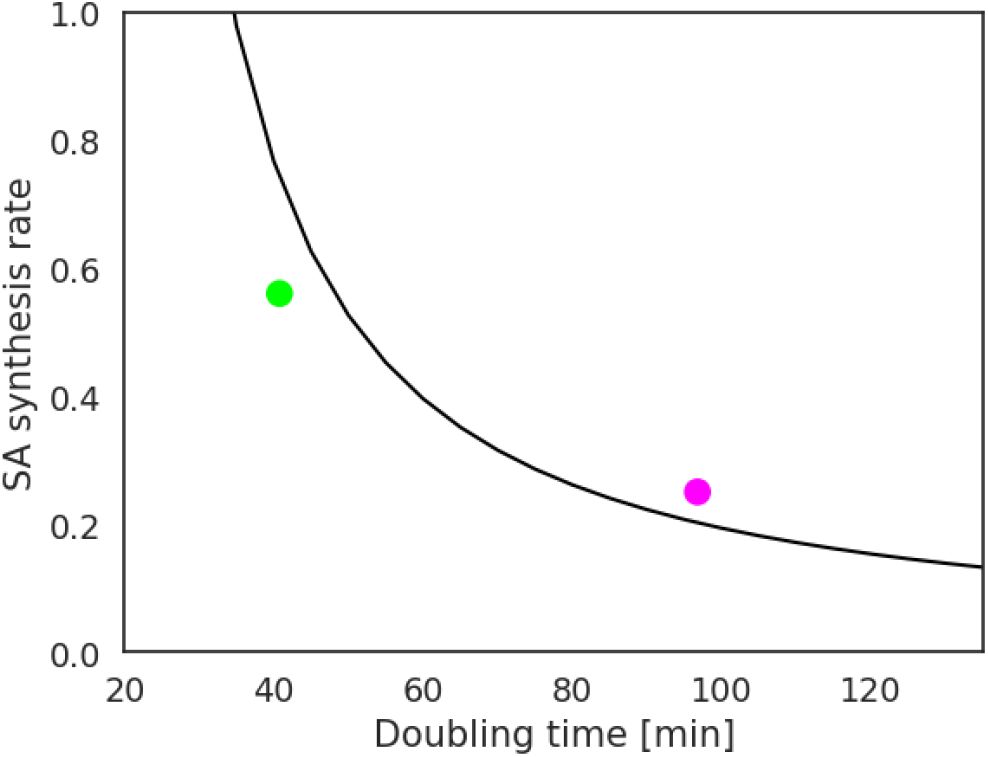
Model prediction for growth-rate dependent susceptibility to mecillinam. The black line shows the model prediction for the threshold surface area synthesis rate (in *μ*m^2^/h) calculated using data from ***Taheri-Araghi et al. (2015)*** for birth volume as a function of doubling time, and the model relation *Y*_t_ = 9*V* ^2*/*3^*/*(4*πT*_0_). The points show our microfluidic measurements of surface area synthesis rate for cells exposed to mecillinam in narrow channels (Fig. 2), averaged over all cells and over the timeframe 13-14h (mecillinam is added at 10h). Lime green: MOPSgluRDM; magenta: MOPSgluMIN. The data point for MOPSgluRDM lies below the predicted threshold line, indicating insufficient surface area synthesis for proliferation in the presence of mecillinam. The data point for MOPSgluMIN lies above the predicted threshold line, indicating that the surface area synthesis rate is sufficient for continued proliferation in the presence of mecillinam.

### Growth-rate threshold for efficacy of mecillinam

Our model points to a growth-rate threshold for the efficacy of mecillinam. To test this, we performed plate-reader growth experiments on 6 media with either glucose or glycerol as carbon source - MOPSgluMIN, MOPSgluCAA, MOPSgluRDM, MOPSglyMIN, MOPSglyCAA, MOPSglyRDM, at pH6.5 and 34°C, with antibiotic-free doubling times, measured in the plate reader, between 33min and 146min (Fig. 4a,b; see Materials and Methods). Bacteria were pre-grown on the relevant media, then exposed to mecillinam at 1.5*μ*g/ml. In the presence of mecillinam at 1.5*μ*g/ml, our plate reader data fell into two categories (Fig. 4c). For richer media (MOPSGluRDM, MOPSGluCAA, MOPSGlyRDM and MOPSGlyCAA), we observed initial growth followed by lysis, consistent with our shake flask experiments for rich media (Fig. 1). On these media, the killing dynamics are faster on media with a shorter antibiotic-free doubling time, consistent with previous work (***Lee et al., 2018)***. In contrast, on poorer media (MOPSgluMIN and MOPSglyMIN), we observed either sustained growth (MOPSgluMIN) or initial growth followed by a later slow decline in OD (MOPSglyMIN). To clarify the growth behaviour on the MOPSglyMIN, we repeated our experiments for MOPSglyMIN and MOPSglyRDM at the somewhat higher pH of 7.5. At the higher pH, we observed sustained growth in the presence of mecillinam for the poorer media, MOPSglyMIN, but lysis on the richer media MOPSglyRDM (Fig. 4c; orange and olive green data respectively). We speculate that an additional, pH-dependent factor may be at play for cells grown at pH6.5 on MOPSglyMIN (such as stress-related filamentation).

**Figure 4.**
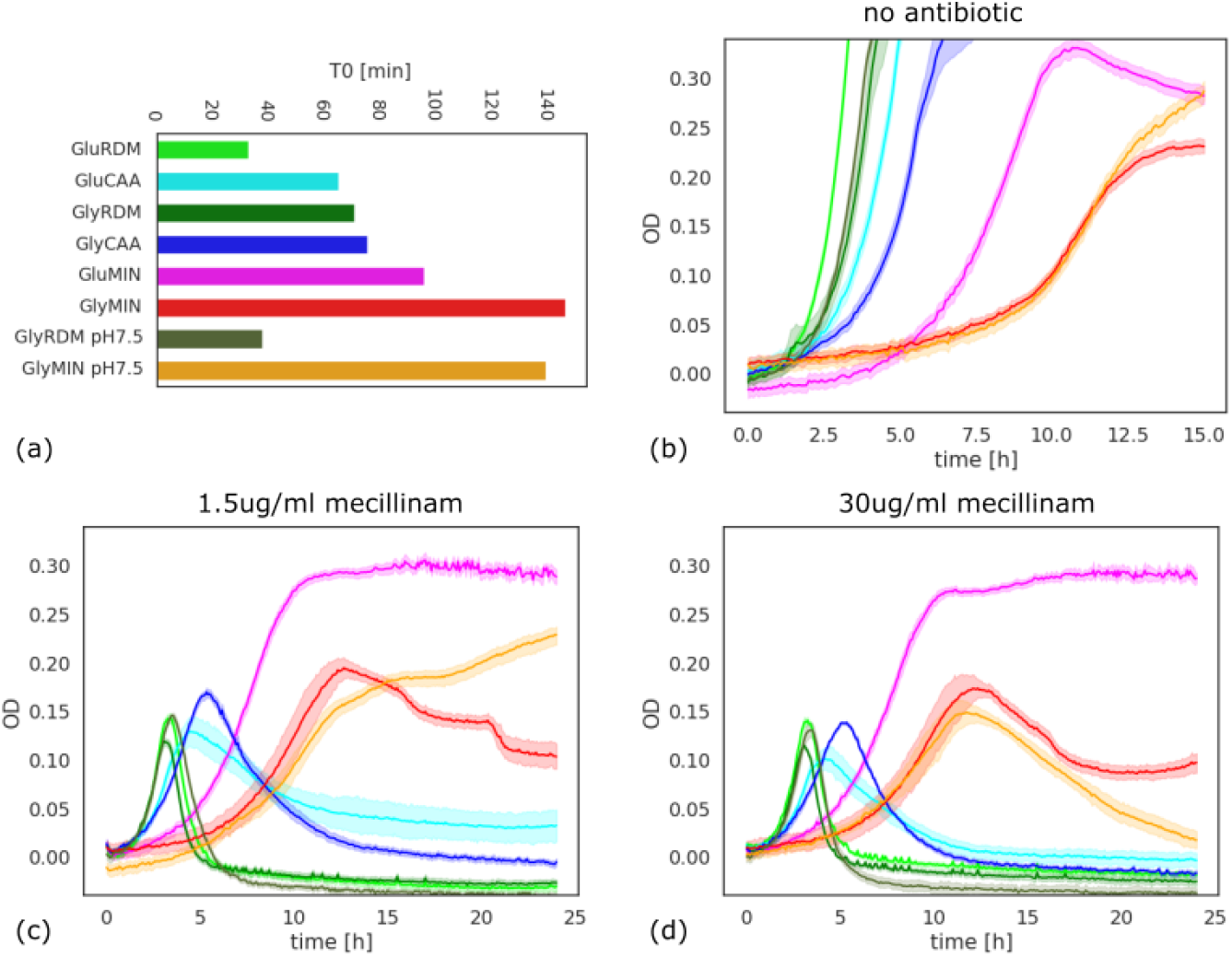
Growth-rate dependence of mecillinam susceptibility. Plate reader growth curves for *E. coli* strain RJA002 measured on a range of MOPS-based growth media. The media (at pH 6.5, 34°C) were MOPSGluRDM (shown in lime-green), MOPSGluCAA (cyan), MOPSGluMIN (magenta) and the equivalent media with glycerol as carbon source: MOPSGlyRDM (mid green), MOPSGlyCAA (blue), MOPSGlyMIN (red). We also performed additional experiments at pH 7.5, 34°C for MOPSGlyRDM (olive green) and MOPSGlyMIN (orange). Further details are given in the Materials and Methods. (a): doubling times for early-time plate reader growth curves, measured in control experiments without antibiotic. (b): Corresponding no-antibiotic growth curves. (c): Growth curves on all media in the presence of mecillinam at 1.5*μ*g/ml. (d) Corresponding data for a higher concentration of mecillinam, 30*μ*g/ml.

Taken together, our plate reader data support the existence of a growth-rate threshold for the efficacy of mecillinam at 1.5*μ*g/ml.

To investigate further, we also performed plate reader experiments at the higher mecillinam concentration of 30*μ*g/ml. Our model predicts that poor media conditions protect cells from mecillinam, even at high concentrations, since proliferation can proceed in the absence of any elongasome activity. Interestingly this appears to be the case for the glucose minimal media (MOPSglu-MIN), where we observed sustained growth even in the presence of 30*μ*g/ml mecillinam. However, this was not the case for the glycerol minimal media (MOPSglyMIN): here the OD declined at later times, at both pH6.5 and pH7.5. Therefore, slow-growing cells can be protected from mecillinam, even at very high doses, but this effect may also be conditional on other physiological factors that are not included in our model.

### Altering cell geometry can protect against mecillinam

Our model suggests that cell morphology plays a central role in growth-rate dependent susceptibility to mecillinam. Therefore we hypothesized that constraining morphology by confining cells in microfluidic channels could change their mecillinam susceptibility. We tested this prediction by analysing the fate of exponentially growing cells on rich and poor media (MOPSgluRDM and MOPSgluMIN) after exposure to mecillinam in the 1.6×1.6*μ*m (width × height) channels of our microfluidic mother machine (Fig. 5a; see Materials and Methods for details). The channels were pre-treated with bovine serine albumin (BSA) to reduce friction (see Supplementary Movies). As predicted, we observed continued growth and division for laterally confined mecillinam-treated cells on rich media, MOPSgluRDM (Fig. 5b and Supplementary Movie 1) – suggesting that confinement protects cells from mecillinam-induced lysis. Cells grown in channels on poor media (MOPSgluMIN) also continued to grow after exposure to mecillinam, as expected (Supplementary Movie 2).

**Figure 5.**
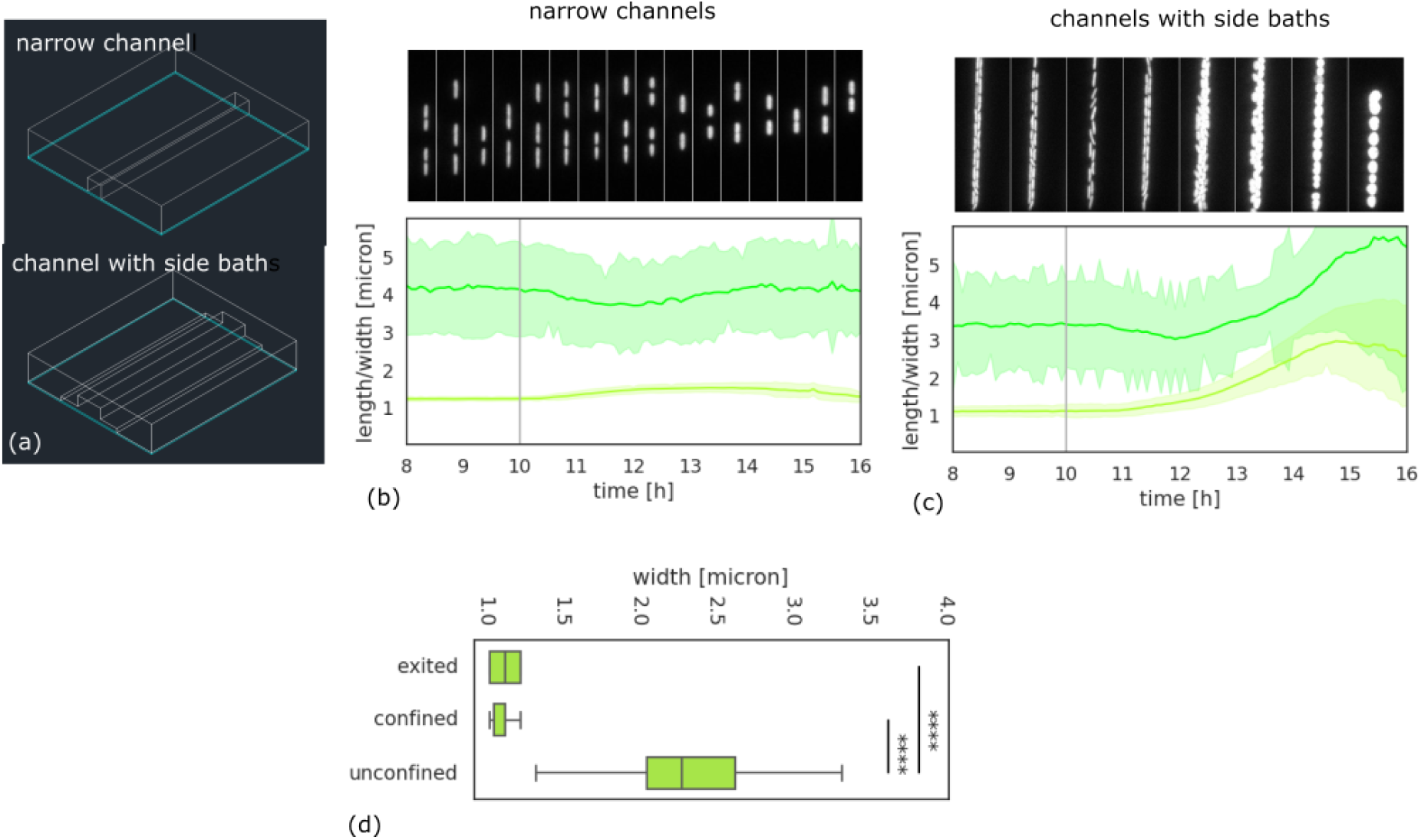
Geometric confinement protects cells from mecillinam. Microfluidic mother machine experiments for *E. coli* strain RJA002 grown on rich media (MOPSGluRDM), before and after addition of mecillinam at 1.5*μ*g/ml (sufficient to cause lysis on this media for cells in bulk, Fig. 1). Mecillinam addition at 10h is indicated by a vertical line. The images show snapshots (corresponding to the time axis of the plots) of a representative channel within the mother machine. (a): Geometry of the narrow channels and channels with side baths. (b) Results for cells grown in narrow channels. The cell morphology alters slightly upon addition of mecillinam, since cells spread laterally to fill the channels, but the cells continue to proliferate. (c): Cells grown in channels with lateral side baths. After mecillinam addition, the cells expand laterally into the side baths, their morphology becomes disordered and they lyse. (d): Cells that were confined in the narrow channels retain their morphology after leaving the channels. The cell width is plotted for cells that have exited the channels, those that are still confined in the channels, and for cells in the same device that remained in the feed channel and were not confined during the experiment. The latter group of cells become large and round, similar to those in bulk cultures.

To verify that the protection of the fast-growing cells from mecillinam was due solely to geometric confinement, we repeated the same experiment (with rich media MOPSgluRDM) using a mother machine with a different channel design in which the 1.4 × 1.4*μ*m channels were flanked by 7 × 0.5*μ*m (width × height) side baths (Fig. 5a). This design enabled the cells to swell laterally into the side baths while still remaining mostly in single or double file within the growth channel. Prior to mecillinam addition, we observed no effect of the side baths on cell growth. However, after addition of mecillinam, the cells swelled to occupy the side baths and rapidly lysed (Fig. 5c and Supplementary Movie 3). Therefore, the absence of lateral confinement restores susceptibility to mecillinam.

We were also curious about the morphology of mecillinam-treated cells after they exited the narrow channel mother machine. Since PG is rigid, we would expect the spherocylindrical morphology that is imposed by geometrical confinement to be maintained for some time after cells exit the channels. To check this, we grew cells for 10 hours on MOPSgluRDM in the narrow channel device, exposed them to mecillinam for a further 3h, then stopped the flow and imaged cells in brightfield as they were leaving the channels (Materials and Methods). Indeed, the rod-like cell morphology was maintained in cells that had existed the channels (Fig. 5d). Our imaging also picked up some cells that had remained outside the channels throughout the experiment (Supplementary Movie 4). As expected, these cells had a significantly different morphology, being round and swollen, similar to the morphologies observed for bulk liquid cultures (Fig. 5d; Fig. 1).

## Discussion

Growth-rate dependent efficacy of *β*-lactam antibiotics has long been known (***Tuomanen et al., 1986; Cozens et al., 1986; Hadas et al., 1995; Lee et al., 2018)***, but mechanistic understanding of this phenomenon has been lacking. In this study, we propose that the growth-rate dependent action of the *β*-lactam antibiotic mecillinam can be explained in simple terms as a perturbation of the balance between the cellular rates of surface area and volume synthesis. Mecillinam binds specifically to the PBP2 TPase component of the elongasome leading to a spherical morphology. While the rate of volume synthesis remains unperturbed by mecillinam, the rate of surface area synthesis decreases, since elongasome activity is compromised. Our data shows that, on rich media, mecillinam treated cells do not maintain a stable size but instead swell over several cell cycles before lysing – in contrast, on poor media, cells can proliferate stably with a round morphology even at mecillinam concentrations well above the EUCAST MIC. Consistent with these observations, a simple geometric model predicts the existence of a “growth-rate threshold” for mecillinam efficacy: stable growth is predicted only for cells growing on media where the antibiotic-free growth rate is below the threshold. Also consistent with this picture, geometric confinement of cells in microfluidic channels can protect them from the action of mecillinam under rich media conditions where unconfined cells would lyse.

Recent work has highlighted the relevance of surface area vs volume homeostatis in cell morphology (***Harris and Theriot, 2016)*** and antibiotic response (***Serbanescu et al., 2020; Ojkic et al., 2022; Cylke et al., 2022)***, although the underlying mechanisms for this homeostatis are only now being revealed (***Shi et al., 2021; Oldewurtel et al., 2021, 2023)***. Our study strengthens the case that surface area vs volume synthesis is an important simple concept that can be useful in understanding mechanistically complex phenomena in microbial physiology.

*β*-lactams account for the majority of clinical antibiotic usage, hence understanding their interplay with bacterial growth physiology is relevant, especially as this may help explain why antibiotic efficacy in the body may be different than that measured under lab conditions (***Zak and Sande, 1982; Cozens et al., 1986)***. In future work, it will be interesting to extend our study to other antibiotics, including both other elongasome-inhibiting agents (e.g. A22, imipenem) and antibiotics that primarily target the divisome (e.g. aztreonam).

Taken together, our work contributes to the emerging understanding of how simple physiological principles can be used to explain complex effects of antibiotics, with the ultimate aim of achieving more effective and efficient antibiotic use.

## Materials and Methods

### Bacterial strains and growth media

*E. coli* strain RJA002 was used in this work. RJA002 is a derivative of the wild-type strain MG1655 that contains a chromosomal copy of the yellow fluorescence protein gene under the control of the constitutive *λ*P_*R*_ promoter, together with a chloramphenicol resistance cassette, as previously described (***Lloyd and Allen, 2015)***.

Our experiments were performed in Neidhardt’s defined MOPS media, consisting of potassium morpholinopropane sulfonate (MOPS) buffer, to which are added essential nutrients (***Neidhardt et al., 1974)***, as well as a carbon source and in some cases, additional amino acids and nucleotides. Minimal media (e.g. MOPSgluMIN) consisted of 10x MOPS base medium (***Brouwers et al., 2020)***, 0.132M K_2_HPO_4_ and a carbon source, added to a final w/v of 2%. In most experiments the carbon source was glucose, but we also performed experiments with glycerol as carbon source. For casamino-acid media (e.g. MOPSgluCAA), the minimal media was supplemented with casamino acids (CAA) to a final concentration of 0.2% (w/v) from a 10% casamino acid solution. Rich media (e.g. MOPSgluRDM) consisted of 10x MOPS base medium, 0.132M K_2_HPO_4_ and additionally 10x ACGU supplement (containing nucleotides) and 5x EZ supplement (containing amino acids) (***Neidhardt et al., 1974)***. The ACGU and EZ supplements were prepared as described by (***Brouwers et al., 2020)***.

The half-life of mecillinam in MOPS media is relatively short under the usual conditions of pH 7.4 and a temperature of 37°C (**?**). Therefore, our experiments were performed at pH 6.5 and 34°C, for which antibiotic has a half-life of more than 6 hours. pH adjustment was achieved by titration of 1M HCl to the growth media.

### Shake flask experiments (Fig. 1)

The shake-flask experiments shown in Fig. 1 were performed using 10ml cultures in 50ml conical tubes. The tubes were inoculated with steady state cultures that had previously been maintained in the exponential growth regime for more than 10-20 generations, as detailed in Supplementary Methods. OD was measured at 600 nm for 1 ml samples using a Cary UV spectrophotometer and polystyrene cuvettes (FisherBrand, FB55146). Colony forming unit (CFU) measurements were performed by performing tenfold serial dilutions up to 10^−8^in PBS buffer and spotting onto LB agar plates which were incubated overnight before counting (for further details see Supplementary Methods).

### Phase contrast microscopy (Fig. 1)

Fig. 1 shows microscopy images of samples taken at different times from our shake flask cultures (see above). Images were taken using an inverted phase contrast microscope (Nikon Ti-U) with a 100×, 1.45 NA phase oil objective in combination with a digital-camera (CoolSNAPHQ2). For imaging, 2 − 4*μ*l of cell culture was sandwiched between a cover slip and microscope slide, and sealed with VALAP (1:1:1 vaseline, lanolin and beeswax).

### HPLC analysis of extracted peptidoglycan (Table 1)

High-performance liquid chromatography (HPLC) was used to analyse PG extracted from 500ml cultures of strain RJA002 after 2h exposure to mecillinam at 1.5 *μ*g/ml. We compared cultures grown on MOPSgluMIN or MOPSgluRDM at pH 6.5 and 34°C, vs no-mecillinam controls. Two replicate experiments were performed for each condition. PG was extracted and processed as described (***Glauner, 1988)***, the muropeptides were released with cellosyl (kindly provided by Hoechst, Frankfurt, Germany), reduced with sodium borohydride and separated by HPLC using the Glauner method (***Glauner, 1988)***. Further details, as well as the full dataset, are given in the Supplementary Methods.

## Microfluidic experiments (Fig. 2 and Fig. 5)

The microfluidic experiments of Fig. 2 and Fig. 5 were performed in a mother machine setup (***Panlilio et al., 2021; Long et al., 2013; Wang et al., 2010)***, in which cells are trapped in narrow channels that have been pre-treated with bovine serine albumin (BSA) to prevent sticking. Medium flow was maintained at a rate of 3*μ*l/min using two syringe pumps; media could be switched (for antibiotic exposure using a Y junction. Cells were grown in the mother machine for at least 10h before the antibiotic was administered. To determine the fate of cells after geometric confinement (Fig. 5c), flow was stopped after 3h exposure to mecillinam and cells were imaged in brightfield as they were leaving the channels.

Two different mother machine designs were used. To test whether lateral confinement enabled bacterial survival, we used a mother machine with simple 25*μ*m long and 1.6*μ*m wide channels. To determine whether death occurred in an equivalent environment without lateral confinement, we deployed a mother machine with a 75*μ*m long growth channel (of height and width 1.4*μ*m), flanked by side baths of 7*μ*m width and 0.5*μ*m height. This enabled cells to swell laterally into the side baths while still remaining mostly in single or double file within the growth channel. The mother machine devices were fabricated from PDMS using standard procedures (as detailed in Supplementary Methods).

The mother machines were imaged using a customised Nikon Eclipse Ti-E microscope in brightfield and epifluorescence using custom scripts, a 40x air objective (Plan Apo *λ* 40 ×, N.A. 0.95, Nikon) an an EMCCD camera (Andor iXon DU-897, Oxford Instruments Industrial Products Ltd.) Images were taken every 5 minutes at 0.1 or 0.5s exposure times for brightfield and fluorescence, respectively and 0 or 300 camera gain. Brightfield illumination was provided by a red LED (LUXEON Z). For YFP imaging we used a blue LED (LUXEON Z), a 495/20nm excitation filter and a 540/30nm emission filter in a filter cube (Nikon) with a 515nm dichroic mirror. To image cells leaving the channel we used a Grasshopper 3 GS3-U3–23S6M-C CMOS camera (FLIR Integrated Imaging Solutions GmbH, Germany) in brightfield with the same illumination settings, at 10 FPS.

For the thin channels, data analysis was performed using Bugpipe in Matlab (***Panlilio et al., 2021)***, followed by python for plotting. Cell volume and surface area were estimated by assuming a spherocylindrical cell shape. Exponential growth rates for volume and surface area were obtained by linear fitting of logarithmic plots, using only cells that appeared in at least three consecutive frames. To analyse images from the channels with side baths we used ilastik (***Berg et al., 2019)*** and SuperSegger (***Stylianidou et al., 2016)***, together with Matlab and Python (see Supplementary Methods). The widths of cells leaving the channels were estimated manually using ImageJ (***Schneider et al., 2012)***.

### Plate reader growth curves (Fig. 4)

The plate reader growth data on multiple growth media (Fig. 4) were obtained as follows. Cultures of strain RJA002 were inoculated from a single colony in 5ml LB medium and grown for 8h at 37°C with 200 rpm shaking. The resulting precultures were diluted 1:100,000 into fresh relevant MOPS media (with pH adjusted to 6.5 where relevant, volume 5ml) and grown overnight at 37°C with 200 rpm shaking. The overnight cultures were diluted again in relevant MOPS media (adjusted to OD (600nm) = 0.01) and grown to the exponential phase (OD (600nm) = 0.09-0.3 depending on fast or slow growth) at 37°C with 200rpm shaking.

The resulting exponential cultures were diluted in relevant MOPS media to OD (600nm) = 0.02 and were used to inoculate a Greiner Bio-One 96 F-bottom Cellstar well plate (KL43.1) which had been pre-filled with the relevant growth media and mecillinam concentrations (100*μ*l per well). The inoculum volume was also 100*μ*l, leading to a starting OD (600nm) in each well of approximately 0.01. The plates were sealed with Breathe-Easy membranes (Merck Z380059) and grown in a ClarioStar plus spectrophotometer at 34°C with 600rpm double orbital shaking for 48h. Growth was monitored by measuring OD (600nm) every 5min. All experiments were performed in at least 4 replicate wells (technical replicates) per plate, for 2 biological replicates.

To process the data, we first subtracted from all curves dynamical background data corresponding to OD (600nm) curves measured for control wells with no inoculum, for the relevant media and antibiotic concentration. The background curves were computed as averages over 8 replicate control wells for each plate and each media-antibiotic combination. Antibiotic-free exponential growth rates for each growth medium, reported in Fig. 4a, were obtained from linear fits of the logarithm of the averaged growth curves (8 replicates) in the early growth regime (generally OD (600nm) between 0.08 and 0.2, although the range was adjusted slightly for some datasets).

## Acknowledgments

The authors acknowledge valuable discussions with Simon Foster, Jamie Hobbs, Meriem el Karoui, Sebastian Jaramillo Rivera, Nikola Ojkic, Abimbola Olulana, Teuta Pilizota, Matt Scott, Bartek Waclaw and Sven van Teeffelen. We also thank Conrad Woldringh and Tanneke den Blaauwen fortheir very helpful advice and assistance in the early stages of this work. This work was supported by an EPSRC DTA studentship to RB, by the European Research Council under Consolidator grant 682237 EVOSTRUC and by EPSRC under grant EP/T002778/1. The work was also supported by a Royal Society University Research Fellowship and by the Excellence Cluster Balance of the Microverse (EXC 2051 Project-ID 390713860) funded by the Deutsche Forschungsgemeinschaft (DFG). For the purpose of open access, the author has applied a Creative Commons Attribution (CC BY) licence to any Author Accepted Manuscript version arising from this submission.

## Notes

### Competing Interest Statement

The authors have declared no competing interest.

