## Supplemental methods and figures for "Surface versus volume synthesis governs growth-dependent efficacy of a *β*-lactam antibiotic"

### Surface versus volume synthesis governs growth-dependent efficacy of a $\beta$-lactam antibiotic: Supplementary Material

**Rebecca Brouwers, Leonardo Mancini, Sharareh Tavaddod, Jacob Biboy, Marco Mauri, Elizabeth Tatham, Marie-Luise Enghardt, Ariane Zander, Pietro Cicuta, Waldemar Vollmer and Rosalind J. Allen**

#### Supplementary Movies

- **Supplementary Movie 1**: Microfluidic mother machine experiment for *E. coli* strain RJA002 grown on rich media (MOPSgluRDM), in narrow BSA coated channels, before and after addition of mecillinam at 1.5$\mu$g/ml (Fig. 5b of the main text). Mecillinam is added at 10h.
- **Supplementary Movie 2**: Mother machine experiment for *E. coli* RJA002 grown on rich media (MOPSgluRDM), in BSA-coated channels with side baths before and after addition of mecillinam at 1.5$\mu$g/ml (Fig. 5c of the main text). Mecillinam is added at 10h.
- **Supplementary Movie 3**: Mother machine experiment for *E. coli* RJA002 grown on poor media (MOPSgluMIN), in narrow BSA coated channels before and after addition of mecillinam at 1.5$\mu$g/ml (Fig. 5c of the main text). Mecillinam is added at 10h.
- **Supplementary Movie 4**: Mother machine experiment in narrow channels on MOPSgluRDM, as for Supplementary Movie 1, but flow is stopped 3h after mecillinam addition and cells are imaged in brightfield at 10 frames per second as they exit the channels.

#### Supplementary Methods

##### Steady state cultures

The shake flask experiments shown in Fig. 1 of the main text were inoculated with cells that had been previously grown to steady state under the relevant conditions. The following protocol was used to prepare a steady state culture (***Grover and Woldringh, 2001***). Single colonies were used to inoculate 5ml seed cultures in Luria-Bertani medium (LB; Fisher-Scientific, made up in-house, pH 7.0), which were grown for 2h at 37°C with 200 rpm shaking. $10\mu$l of the seed culture was then used to inoculate 5 ml liquid cultures in the chosen growth medium, which were allowed to grow overnight at 37°C with 200 rpm shaking. The overnight cultures were used to inoculate new liquid cultures in the chosen growth medium, which were then maintained in the exponential growth regime for 10–20 generations by diluting periodically in pre-warmed growth medium. The maximum optical density of a cell culture before each dilution was 0.4 at $450$ nm.

##### Colony forming units

For our CFU measurements, shown in Fig. 1 of the main text, the following method was used (***Herigstad et al., 2001***). The *E. coli* culture of interest was vortexed for 6 seconds before an initial 1 ml sample was extracted and serially diluted ten fold up to $10^{-8}$ in PBS (8x1.5 ml eppendorfs had been prepared containing $900\mu$l PBS to which $100\mu$l sample was added). These eight dilutions were pipetted in 6-8 $10\mu$l drops onto LB agar plates as shown in Supplementary Fig. 1. To obtain the time course CFU data of Fig. 1 of the main text, the procedure was repeated for samples taken from the culture of interest at hourly or two hourly intervals. Once the drops had dried on the LB agar plates, the plates were inverted and incubated at 30°C overnight. Viable colony forming units were

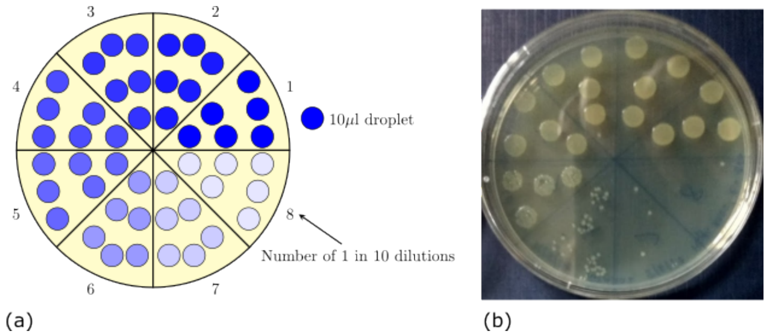


**Supplementary Figure 1. CFU measurement technique** (a): A typical plate set-up with eight segments each containing six $10\mu$l droplets. Each segment is for a different culture dilution with the number referring to the number of 1 in 10 dilutions in PBS. For example segment 7 contains droplets of the sample culture diluted 1 in 10000000. The shade of blue in the droplets is a visual guide to demonstrate the increasing diluted nature of the sample across the segments. (b): Example CFU plate from an experiment with the segments aligned with panel (a).

---

counted the following morning from whichever dilution had 3-30 colonies per $10\mu$l drop (***Herigstad et al., 2001***). This number was then scaled by the corresponding dilution and drop volume to give a measure of the number of colony forming units in 1 ml of culture.

#### Minimum inhibitory concentration (MIC) measurement

The MIC value was measured for the parent strain *E. coli* MG1655 on LB medium. MIC measurements were performed in a 96 well plate, such that the final volume in each well was $200\mu$l. To prepare the antibiotic dilutions, $190\mu$l of growth media was first added to every row, then an additional $150\mu$l was added to the top row giving a total of $340\mu$l. $40\mu$l of antibiotic was the added to the top row to give a total volume of $380\mu$l and a starting concentration of four times the MIC. This was mixed using the electronic pipette by pipetting $190\mu$l up and down, five times. Once mixed $190\mu$l of the top row was pipetted up and added to the next row down, where mixing was again carried out. This resulted in antibiotic concentrations that decreased two-fold down the columns. No antibiotic was added to the bottom row (***CLSI, 2013***; ***Jepson, 2014***). After the antibiotic had been distributed, each well contained $190\mu$l, to which $10\mu$l of *E. coli* MG1655 culture at optical density 0.2 was added, giving a starting inoculation density of optical density 0.01. The 96-well plate was then incubated at 37°C in the plate reader for 24 hours. The MIC was determined as the concentration at which there was no visible growth after the 24 hour incubation; this was 1.5 $\mu$g/ml.

#### HPLC analysis of extracted peptidoglycan

For pre-culture, cells were inoculated in 20 ml MOPSgluRDM and grown at 37°C overnight, washed and added to the relevant media at 34°C with a starting OD (600nm) of approximately 0.05. Mecillinam was added to growing cell cultures and after 2h exposure, lysates were prepared by re-suspending cells in MOPSgluMIN (since mecillinam-exposed cells lyse rapidly if re-suspended in water) and adding them to boiling 8% SDS. PG was prepared as previously described (***Glauner, 1988***). Each aliquot of PG was then digested overnight with cellosyl before the reaction was stopped by heating at 100°C for 10 min. Samples were centrifuged, the supernatant collected and the muropeptides were reduced using sodium borohydride. HPLC analysis was then performed using the Glauner method (***Glauner, 1988***). Representative muropeptide profiles are shown in Supplementary Fig. 2.

#### Microfluidic experiments

##### Bacterial growth conditions

Single colonies of *E. coli* strain RJA002 were inoculated in M63 medium (13.6 g/l $KH_2PO_4$, 0.5 mg/l

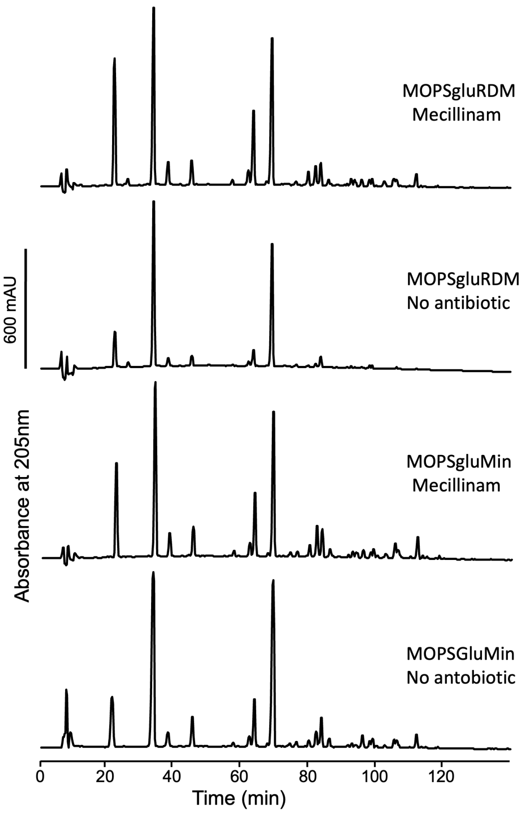


**Supplementary Figure 2. HPLC: muropeptide profiles** Representative muropeptide profiles of RJA002 cells grown in MOPSGluRDM or MOPSGluMIN after 2h in the presence of 1.5$\mu$g/ml mecillinam or without antibiotic. The PG was isolated from cells and the muropeptides were released with cellosyl, reduced with sodium borohydride and separated by HPLC.

---

$FeSO_4 \cdot 7H_2O$, 0.5 g/l $MgSO_4 \cdot 7H_2O$, 1.27 mg/l Thiamine, 2.64 g/l $(NH_4)2SO_4$ and 0.5% w/v Glucose, final pH: 7.2) and grown overnight in conical flasks at 37°C with shaking at 220 rpm. The culture was then concentrated 20 times using centrifugation at 8000g for 2 minutes and loaded for up to 1 hour into mother machines that had been pre-treated with 20mg/ml of BSA (Sigma) for 20 minutes at room temperature. Bacteria in excess were washed off the feeding channel using either fresh MOPSgluRDM or MOPSgluMIN. During imaging, medium flow was provided by two syringe pumps (kdScientific) at a rate of 3 $\mu$l/min. Switches between fresh media and fresh media supplemented with mecillinam were performed through a "Y" junction (Darwin microfluidics). The temperature of the mother machine was kept stable at 34°C by sticking the edges of the glass coverslip onto a custom-built thermoelectric chamber. We note that the syringes containing media were left at room temperature throughout the experiment to preserve mecillinam efficacy, while tubing was woven through tiny holes of the chamber to ensure the medium reaching the cells was already at the desired temperature. Cells were grown in the mother machine for 10h before the antibiotic was administered. To check whether cells leaving the channels would maintain the shape they had within the channel (Fig. 5 of the main text), we performed the experiment as before, flowing antibiotic-free MOPSgluRDM for 10 hours and then switching to medium supplemented with the antibiotic. After 3 hours in the presence of the antibiotic, flow was stopped and cells were imaged in brightfield as they were leaving the channels.

**Microfluidics fabrication** Two different mother machine designs were used. To test whether lateral mechanical confinement of the cell wall enabled bacterial survival, we used a mother machine with simple 25$\mu$m long and 1.6$\mu$m wide channels (Fig). To assay whether death occurred in larger channels, we deployed a mother machine with a 75$\mu$m long, 1.4$\mu$m wide growth channel and 7$\mu$m wide side baths (Fig. 5 of the main text). The height of the growth channel was equal to its width, while the side baths were only 0.5$\mu$m high. This enabled cell swelling while maintaining cells mostly in single or double files. This is important because cells in multiple files can fill the side baths, prompt confinement and limit cell death. PDMS (1 to 10 curing agent to PDMS, Dow Corning) was mixed and degassed and then cast onto mother machine epoxy moulds. After >2h incubation at 60°C, the chips were peeled from the moulds, cut with a scalpel and the inlets and outlets were pierced using a 0.75mm puncher (Rapid-core). The chips were then bond onto glass coverslips (Menzel-Gläser) using a plasma cleaner after a 1min plasma treatment. The bond microfluidics were then incubated for further 10 minutes at 60°C. To connect inlets and outlets to Tygon tubing (Cole Parmer; 0.020" × 0.060" outside diameter), we used 90°bent connectors (Darwin microfluidics, stainless steel, 0.89-0.58 OD-ID, 20 gauge). Media switches (between media without and with mecillinam) were performed through a "Y" connector (Darwin microfluidics). The tubing was then connected to 5 ml plastic syringes (BD plastipak) via blunt needles (Intertronics; stainless steel, straight blunt, 1/2", 23 gauge). Media flow was prompted by syringe pumps (kdScientific). The piece of tubing connecting the syringe containing mecillinam with the Y-junction was filled with antibiotic-less medium before the experiment to guarantee that no mecillinam would we be perfused before the switch.

**Imaging in microfluidics** Mother machines were automatically imaged in brightfield and epifluorescence using custom scripts in a customised Nikon Eclipse Ti-E microscope. Images were taken through a 40x air objective (Plan Apo $\lambda$ 40 ×, N.A. 0.95, Nikon) with an EMCCD camera (Andor iXon DU-897, Oxford Instruments Industrial Products Ltd.) every 5min at 0.1 or 0.5s exposure times for brightfield and fluorescence, respectively and 0 or 300 camera gain. Brightfield illumination was provided by a red LED (LUXEON Z). To observe YFP fluorescence, illumination was provided by a blue LED (LUXEON Z) through a 495/20nm excitation filter and light was collected at a 540/30nm emission in a filter cube (Nikon) mounted with a 515nm dichroic mirror. To monitor whether cells leaving the pistons of the thin channels mother machine would maintain their shape, we took advantage of the larger field of view of a Grasshopper 3 GS3-U3–23S6M-C CMOS camera (FLIR Integrated Imaging Solutions GmbH, Germany) imaging in brightfield with the same illumination settings explained before, at 10 FPS.

**Data analysis** Images from thin channels (Fig. 5 of the main text) were analysed using Bugpipe in Matlab as previously (***Panlilio et al., 2021***). The extracted data was then further handled in Python. Because images only contain length, width, time and fluorescence information, volume and surface of the cells were estimated by modelling cells as cylinders capped by two hemispheres (where cell width = sphere diameter). Growth rates of volume and surface were only estimated for cells that appeared in at least three consecutive frames. Exponential growth rates were extracted by taking the natural logarithm of the quantities (volume or surface) and fitting straight lines through them. The slope of such lines is our growth rate. In the channels with side baths (Fig. 5 of the main text) cells treated with mecillinam take up less canonical shapes that we could not segment through Bugpipe. For these samples we used ilastik (***Berg et al., 2019***) to produce cell masks, separating foreground from background. To refine cell segmentation and extract length and width of these cells, we used Supersegger (***Stylianidou et al., 2016***), Matlab and Python. Because Supersegger appeared to segment best for images in phase contrast (which have greyvalues of the background > greyvalues of the foreground), we uniformly set the background to an arbitrary value of 40000, essentially mimicking the ratios appearing in phase contrast images. The images thus generated were segmented through Supersegger and the output data handled in Python. The scripts for Bugpipe, ilastik and Supersegger are available (***Berg et al., 2019***; ***Stylianidou et al., 2016***). In the channels with side baths, we note that cells can at times be in double files and very close together. Therefore the threshold between background and foreground used when training ilastik has to be slightly higher than the one used by bugpipe. Because of this, cell size tends to be slightly underestimated during segmentation (and roughly by 1 pixel on each end) in the large channels. We correct for this by adding 2 pixels to all of the measurements. Estimation of the widths of cells leaving the channels (Fig. 5 of the main text) was performed manually using ImageJ (***Schneider et al., 2012***).

#### Supplementary Data

| Muropeptide | Relative peak area (%) in PG from RJA002 cells grown in MOPSgluRDM | MOPSgluMIN | MOPSgluRDM | MOPSgluMIN |
|---|---|---|---|---|
| | No antibiotic | | Mecillinam | |
| **Tri** | 9.2 ± 0.5 | 9.0 ± 0.3 | 15.1 ± 0.5 | 13.0 ± 0.6 |
| **TetraGly4** | 1.1 ± 0.0 | 0.0 | 0.8 ± 0.0 | 0.0 |
| **Tetra** | 41.6 ± 1.6 | 35.7 ± 1.7 | 25.0 ± 0.5 | 24.8 ± 1.2 |
| **PentaGly5** | 0.0 | 0.0 | 0.0 | 0.0 |
| **Di** | 2.0 ± 0.1 | 2.2 ± 0.1 | 3.1 ± 0.1 | 3.2 ± 0.0 |
| **Penta** | 0.0 | 0.0 | 0.0 | 0.0 |
| **Tri-LysArg** | 3.9 ± 1.9 | 3.6 ± 0.1 | 3.9 ± 1.0 | 4.0 ± 0.1 |
| **TetraTri(DAP)Gly4** | 0.0 | 0.0 | 0.0 | 0.0 |
| **TriTri(DAP)** | 0.0 | 0.4 ± 0.1 | 0.8 ± 0.3 | 0.7 ± 0.0 |
| **TetraTri(DAP)** | 1.4 ± 0.3 | 1.4 ± 0.1 | 1.7 ± 0.5 | 1.9 ± 0.0 |
| **TetraTetraGly4** | 0.0 | 0.1 ± 0.1 | 0.0 | 0.0 |
| **TetraTri** | 3.9 ± 1.0 | 5.2 ± 0.2 | 10.5 ± 0.1 | 8.2 ± 0.1 |
| **TetraPentaGly4** | 0.2 ± 0.2 | 0.1 ± 0.2 | 0.6 ± 0.0 | 0.4 ± 0.0 |
| **TetraTetra** | 30.1 ± 1.0 | 28.1 ± 1.0 | 21.2 ± 0.3 | 20.7 ± 0.6 |
| **TetraAnh** | 0.0 | 0.3 ± 0.1 | 0.2 ± 0.3 | 0.5 ± 0.1 |
| **TetraPenta** | 0.5 ± 0.1 | 0.7 ± 0.0 | 0.5± 0.0 | 0.8± 0.0 |
| **TetraTetraTri** | 0.1 ± 0.2 | 0.7 ± 0.0 | 1.8 ± 0.1 | 1.4± 0.0 |
| **TetraTri LysArg** | 1.8 ± 1.2 | 1.9 ± 0.4 | 3.8 ± 1.8 | 4.0 ± 0.2 |
| **TetraTetraTetra** | 2.4 ± 0.1 | 3.3 ± 0.4 | 2.7 ± 0.2 | 3.6 ± 0.1 |
| **TetraTriAnh I** | 0.0 | 0.3 | 0.9 ± 0.1 | 0.7± 0.0 |
| **TeraTriAnh II** | 0.0 | 1.0 ± 0.6 | 0.7 ± 0.1 | 0.7± 0.0 |
| **TetraTetraAnh I** | 0.6 ± 0.1 | 0.7 ± 0.0 | 0.7 ± 0.1 | 0.9 ± 0.2 |
| **TeraTetraAnh II** | 0.7 ± 0.1 | 0.9 ± 0.0 | 0.8 ± 0.1 | 1.1± 0.0 |
| **TetraTetraTriAnh** | 0.0 | 0.7 ± 0.4 | 1.0 ± 0.1 | 1.5 ± 0.8 |
| **TetraTetraTetra Anh** | 0.2± 0.3 | 1.3 ± 0.4 | 1.1 ± 0.4 | 1.8 ± 0.2 |
| **TetraTetraTetraTetra Anh** | 0.0 | 1.0 ± 0.6 | 0.8 ± 1.1 | 2.1 ± 1.0 |
| **all known** | 99.7 ± 0.5 | 98.7 ± 1.4 | 97.6 ± 0.9 | 96.0 ± 0.8 |

**Supplementary Table 1.** Muropeptide composition of RJA002 grown in MOPSgluRDM and MOPSgluMIN media after exposure (or not) to 1.5$\mu$g/ml mecillinam for 2 h.

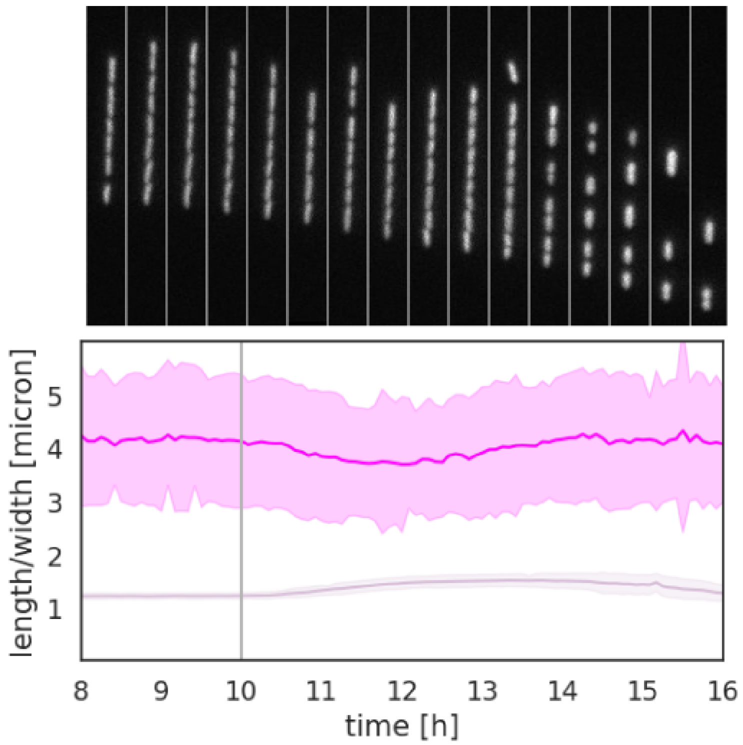


**Supplementary Figure 3. Microfluidic experiment on poor media** Microfluidic mother machine experiments for *E. coli* strain RJA002 grown on poor media (MOPSGluMIN), before and after addition of mecillinam at 1.5$\mu$g/ml. Mecillinam addition at 10h is indicated by a vertical line. The images show snapshots (corresponding to the time axis of the plots) of a representative channel within the mother machine.
